# Dopamine manipulations modulate paranoid social inferences in healthy people

**DOI:** 10.1101/2019.12.18.874255

**Authors:** J.M. Barnby, V. Bell, Q. Deeley, M.A. Mehta

## Abstract

Altered dopamine transmission is thought to influence the formation of persecutory delusions. However, despite extensive evidence from clinical studies there is little experimental evidence on how modulating the dopamine system changes social attributions related to paranoia, and the salience of beliefs more generally. 27 healthy male participants received 150mg L-DOPA, 3mg haloperidol, or placebo in a double blind, randomised, placebo-controlled study, over three within-subject sessions. Participants completed a multi-round Dictator Game modified to measure social attributions, and a measure of belief salience spanning themes of politics, religion, science, morality, and the paranormal. We preregistered predictions that altering dopamine function would affect i) attributions of harmful intent and ii) salience of paranormal beliefs. As predicted, haloperidol reduced attributions of harmful intent across all conditions compared to placebo. L-DOPA reduced attributions of harmful intent in fair conditions compared to placebo. Unexpectedly, haloperidol increased attributions of self-interest for opponents’ decisions. There was no change in belief salience within any theme. These results could not be explained by scepticism or subjective mood. Our findings demonstrate the selective involvement of dopamine in social inferences related to paranoia in healthy individuals.

## 1.0 Introduction

Paranoia involves unfounded beliefs that others intend harm (Freeman & Garety, 2014). Epidemiological evidence suggests that paranoia exists on a spectrum in the general population, ranging from mild social concerns to persecutory delusions (Bebbington et al., 2013; Bell & O’Driscoll et al., 2018). Observational and experimental research has identified a range of personal and interpersonal factors that influence paranoia. On the personal level, worry (Startup et al., 2007), insomnia (Freeman et al., 2012) belief inflexibility (Bronstein et al., 2019), and safety behaviours (Freeman et al., 2007) all contribute to the formation and / or maintenance of paranoia. In terms of social factors, social disadvantage and victimisation (Wickham et al., 2014), trauma (Crush et al., 2018), and poor social support (Freeman et al., 2011) all play a role.

Neurobiologically, the subcortical dopamine system has been cited as a candidate for a ‘final common pathway’ on which accumulated biological, psychological and social stresses might have their most significant impact leading to the symptoms of psychosis (Howes & Kapur, 2009; Howes & Murray, 2014) of which persecutory delusions are the most common symptom (Andrade & Wang, 2012). Although the status of subcortical dopamine as a common pathway has been debated (McCutcheon et al., 2019), there remains extensive evidence for the dysregulation of the subcortical dopamine system in psychosis and the paranoia spectrum. Observational PET neuroimaging has found increased striatal dopamine in people at high-risk of progression to psychosis, (Egerton, 2013; Howes et al., 2011) as well as prior to (Howes et al., 2011) and during (Fusar-Poli & Meyer-Lindenberg, 2012) episodes of psychosis. Antipsychotic medication primarily has its effect through antagonism at D_2_ dopamine receptors in the mesolimbic & nigrostriatal pathway (Kapur et al., 2004). Additionally, stimulant drugs which increase activity at mesolimbic D_2_ dopamine receptors raise the risk of psychosis – with over 40% of recreational methamphetamine users developing psychosis (Lecomte et al., 2018) of which paranoid delusions are the dominant symptom (Voce et al., 2019).

The mechanisms that connect dysregulated dopamine to the symptoms of psychosis have been much debated. Several theories have suggested that striatal dopamine is involved in a process of aberrant salience attribution whereby meaningful connections are made between unrelated events or information which form the basis for delusional beliefs (Seeman, 1987; Spitzer, 1995; Kapur et al., 2005). This has been interpreted in terms of the neuromodulatory effect of dopamine on the integration of prediction error in hierarchical Bayesian models of perceptual learning (Corlett et al., 2009; Sterzer et al., 2018). In these models it has been proposed that altered dopamine transmission leads to abnormally strong weighting of perceptual prediction errors that disrupts learning and eventually manifests as delusions. More specifically, recent computational modelling (Diaconescu et al., 2019) and integrative socio-developmental cognitive accounts (Howes & Murray, 2014) have suggested that disruption to dopamine-mediated processes underlying social interaction may be an important explanatory factor in persecutory delusions.

The evidence base for current theories of delusion largely rely on clinical studies, and there are far fewer studies that have taken the additional step of experimentally altering dopamine function in healthy participants to look for causal effects on psychosis-congruent beliefs. Studies have tested the effect of manipulating the dopamine system on the valuation of harm to others (Crockett et al., 2015), self-interest in economic decision-making (Pedroni et al., 2014) and learning about others’ prosociality (Eisenegger et al., 2013). As far as we are aware, no studies to date have tested the effect of altering dopamine function on attributions of others’ intent to harm, the core social attributional process of paranoia (Freeman & Garety, 2014). Similarly, of the few existing pharmacological studies on delusion-related belief mechanisms, Krummenacher et al (2010) found the effect of levodopa on perceptual sensitivity differed depending on levels of paranormal belief, chosen as a non-clinical analogue of delusional ideation. Mohr et al. (2005), also using levodopa, found that laterality of lexical decision processing altered as a function of magical ideation. However, belief salience (Barnby et al., 2019a) has yet to be tested.

Given the importance of experimental pharmacological intervention studies to understand the mechanisms of psychopathology (Tsou, 2012), this study extends this work by examining how modulating dopamine affects i) attributions of harmful intent – a core interpersonal process of paranoia; and ii) salience of paranormal belief – chosen as a non-clinical analogue of delusional ideation and measured alongside salience of other beliefs. Healthy participants took part in a double-blind, within-subjects, randomised placebo-controlled trial of two drugs that alter the dopamine system –L-3,4-dihydroxyphenylalanine (levodopa or L-DOPA) to potentiate presynaptic dopamine, and haloperidol, to primarily block postsynaptic dopamine transmission via D_2_ receptors. At each stage, participants completed a game theoretic social inference task (multi-round Dictator Game; Barnby et al 2019a) where participants were required to attribute the intentions of their partner after their partner had made a monetary decisions, and a measure of belief salience, that included paranormal beliefs (Barnby et al., 2019b).

Given the role of dopamine in paranoia and paranoid delusions, we predicted that haloperidol would reduce attributions of harmful intent and salience of paranormal beliefs based on the observation that dopamine antagonism is the primary therapeutic mechanism of antipsychotics in the treatment of psychosis (Kaar et al., 2019). We predicted that potentiation of dopamine transmission using L-DOPA in healthy participants would increase attributions of harmful intent and the salience of paranormal beliefs, given increased presynaptic dopamine in those at risk of psychosis (Howes & Kapur, 2009). Following Barnby et al. (2019a), we also predicted that haloperidol and L-DOPA would respectively reduce and increase the amount of trials taken to reach a peak level of high harmful intent attribution but not self-interest attributions. All analysis scripts and open data are available on the Open Science Framework (https://osf.io/mr63j/).

## 2.0 Results

The study (Clinical Trials.gov Identifier: NCT03754062) also included the Salience Attribution Task (Esslinger et al., 2012), although data from this task is not reported here. We preregistered the hypotheses and analysis for the multi-round Dictator Game (https://aspredicted.org/6zg2w.pdf) and belief salience measures (https://aspredicted.org/fh495.pdf) prior to unblinding.

We recruited 30 participants in total for the full experimental procedure and kept 27 for analysis. Two participants were removed from the analysis for having incomplete session data. One participant was removed from the analysis for having a very high Green et al Paranoid Thoughts Scale (GPTS; Green et al., 2008) score (104) over two standard deviations away from the mean of the rest of the sample (46.52), potentially making our analysis less conservative.

### 2.1 Demographics and baseline psychometrics

At baseline individuals recorded their age, ethnicity, political orientation, and filled out the Big-5 personality questionnaire (John & Srivastava, 1999), brief O-LIFE schizotypy questionnaire (Mason & Claridge, 2006), Bond and Lader mood rating scale (Bond and Lader, 1974) for each drug condition pre and post dosing, and the Green Paranoid Thoughts Scale. Table 1 describes the distribution of these measures across the sample. Heart rate and blood pressure of participants at baseline and each study day are presented in Table 2.

**Table 1.**
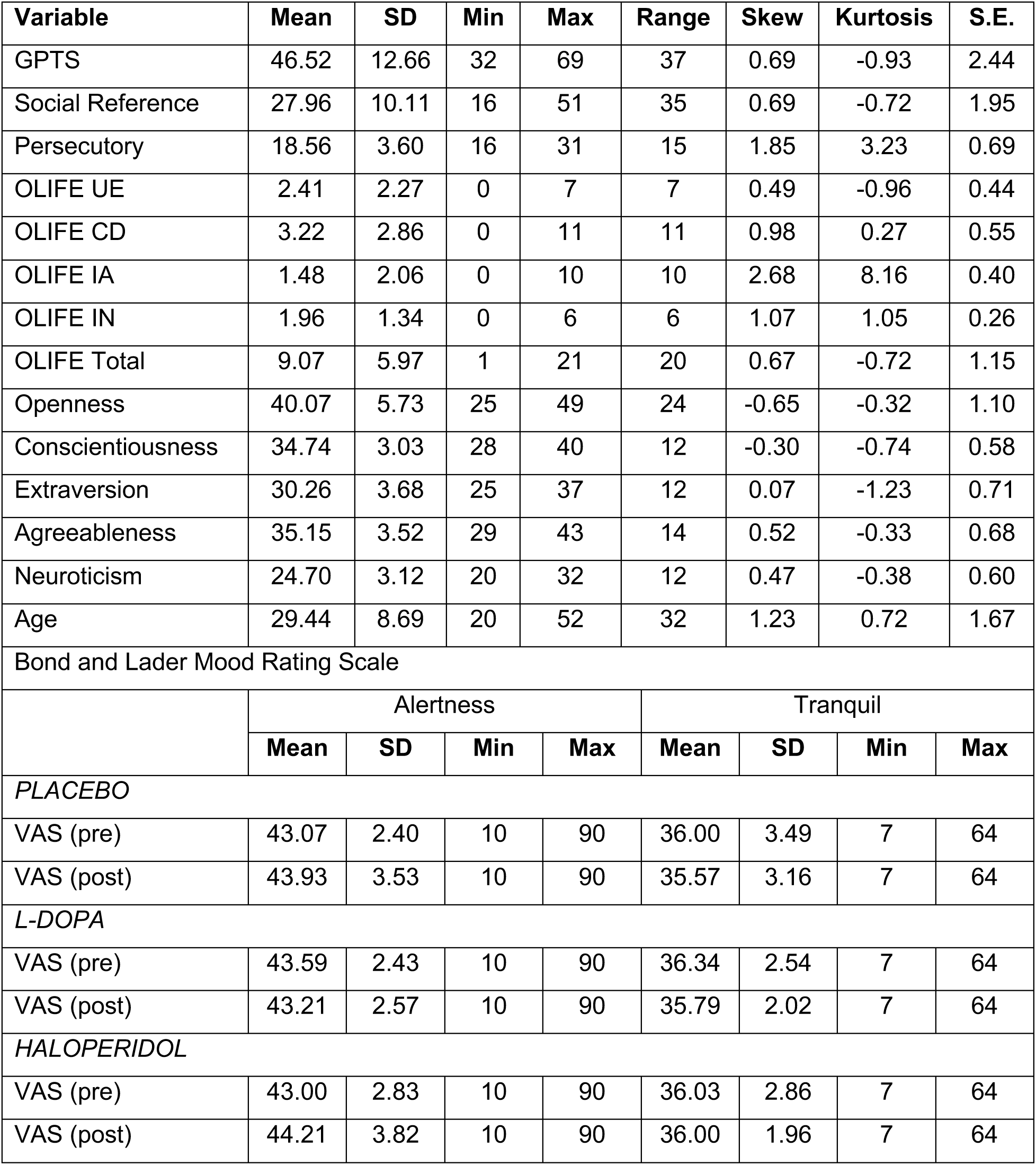
Age, mood ratings, and psychometrics of the included sample (n = 27). Only the Bond and Lader scale (Visual Analogue Scale; VAS) was administered at baseline and subsequent study days, both before and after dosing. OLIFE = Oxford-Liverpool Inventory of Feelings and Experiences (Mason & Claridge, 2006); UE = Unusual Experiences subscale; CD = Cognitive Disorganisation subscale; IA = Introvertive Anhedonia subscale; IN = Impulsive Non-Conformity Subscale.

**Table 2.** Heart rate and blood pressure of participants at baseline and each study day. Formula for differences between sessions are “lmer (Systole/Diastole/HeartRate) ∼ Drug Session + (1|ID)”. Paired t-tests were run for within-session heart rate.

| Condition |  | Systole | Diastole | Heart Rate | Systole | Diastole | Heart Rate | Heart Rate P-val. |
| --- | --- | --- | --- | --- | --- | --- | --- | --- |
| BASELINE | Mean | 123.93 | 71.93 | 66.32 |  |  |  |  |
|  | SD | 9.74 | 9.66 | 10.73 |  |  |  |  |
|  |  | PREDOSE |  |  | POSTDOSE |  |  |  |
| PLACEBO | Mean | 121.39 | 70.82 | 67.86 | 116.25 | 69.75 | 62.14 | 0.11 |
|  | SD | 8.71 | 9.83 | 8.19 | 24.89 | 17.49 | 16.11 |  |
| L-DOPA | Mean | 120.82 | 68.89 | 66.61 | 119.46 | 68.71 | 63.61 | 0.32 |
|  | SD | 9.66 | 7.92 | 10.62 | 9.12 | 9.34 | 9.80 |  |
| HALOPERIDOL | Mean | 121.36 | 69.54 | 67.71 | 116.36 | 67.61 | 60.75 | <0.001 |
|  | SD | 9.78 | 7.95 | 10.03 | 9.68 | 7.77 | 10.09 |  |
| P-val. |  | 0.21 | 0.36 | 0.72 | 0.012 | 0.07 | 0.2 |  |

### 2.2 Multi-round Dictator Game Prediction 1

The multi-round Dictator game was a modified version of the dictator game, where participants were passive receivers of either unfair (100:0) or fair (50:50) splits of money over six trials with three different partners and were only required to infer the intentions of a partner, following each decision, down two dimensions: harmful intent or self interest on a scale of 0-100. While dictators were preprogrammed to either to unfair (always take the money), fair (always split the money) or partially fair (split the money half the time), unlike reinforcement learning paradigms, participants were not required to guess the type of Dictator they were matched with, and their attributions didn’t affect their monetary outcomes in subsequent trials. More details can be found in the methods. In placebo conditions, harmful intent and self interest attributions weren’t correlated with eachother overall or in each Dictator condition (ps > 0.05).

Dopamine manipulation will moderate harmful intent attributions but not self-interest attributions.

We conducted three preregistered analyses for each dictator type. All reported statistics are beta-coefficients of the top model following model averaging unless otherwise stated. See the Supplementary Materials for the mean values collapsed across conditions for harmful intent and self-interest attributions for each drug.

For unfair dictators, compared to placebo, haloperidol reduced harmful intent attributions (−1.155, 95% CI: −1.467, −0.845) but L-DOPA did not (−0.118, 95% CI: - 0.410, 0.169). Compared to haloperidol, L-DOPA increased harmful intent attributions (1.037, 95% CI: 0.736, 1.348). Compared to placebo, haloperidol also increased self-interest attributions (0.650, 95% CI: 0.649, 0.651), but L-DOPA reduced self-interest attributions (−0.021, 95% CI: −0.022, −0.020). Compared to haloperidol, L-DOPA reduced self-interest attributions (−0.670, 95% CI: −0.671, - 0.670).

For partially fair dictators, compared to placebo, haloperidol reduced harmful intent attributions (−0.420, 95% CI: −0.707, −0.133), but L-DOPA did not (0.169, 95% CI: - 0.109, 0.446). Compared to haloperidol, L-DOPA increased harmful intent attributions (0.589, 95% CI: 0.303, 0.874). Compared to placebo, haloperidol also increased self-interest attributions (0.610, 95% CI: 0.362, 0.858) but L-DOPA did not (−0.054, 95% CI: −0.297, 0.188). Compared to haloperidol, L-DOPA reduced self-interest attributions (−0.665, 95% CI: −0.913, −0.416).

For fair dictators, compared to placebo, haloperidol reduced harmful intent attributions (−1.202, 95% CI: −1.202, −1.201), as did L-DOPA (−1.033, 95% CI: −1.034, −1.033). Compared to haloperidol, L-DOPA did not increase harmful intent attributions (0.167, 95% CI: −0.227, 0.561). Compared to placebo, haloperidol did not affect self-interest attributions (0.194, 95% CI: −0.078, 0.469), but L-DOPA decreased them (−0.331, 95% CI: −0.591, −0.070). Compared to haloperidol, L-DOPA reduced self-interest attributions (−0.526, 95% CI: −0.800, −0.254).

Figure 1 illustrates changes to harmful intent attributions and self-interest attributions for each trial, fair and unfair dictators, and drug condition.

**Figure 1:**
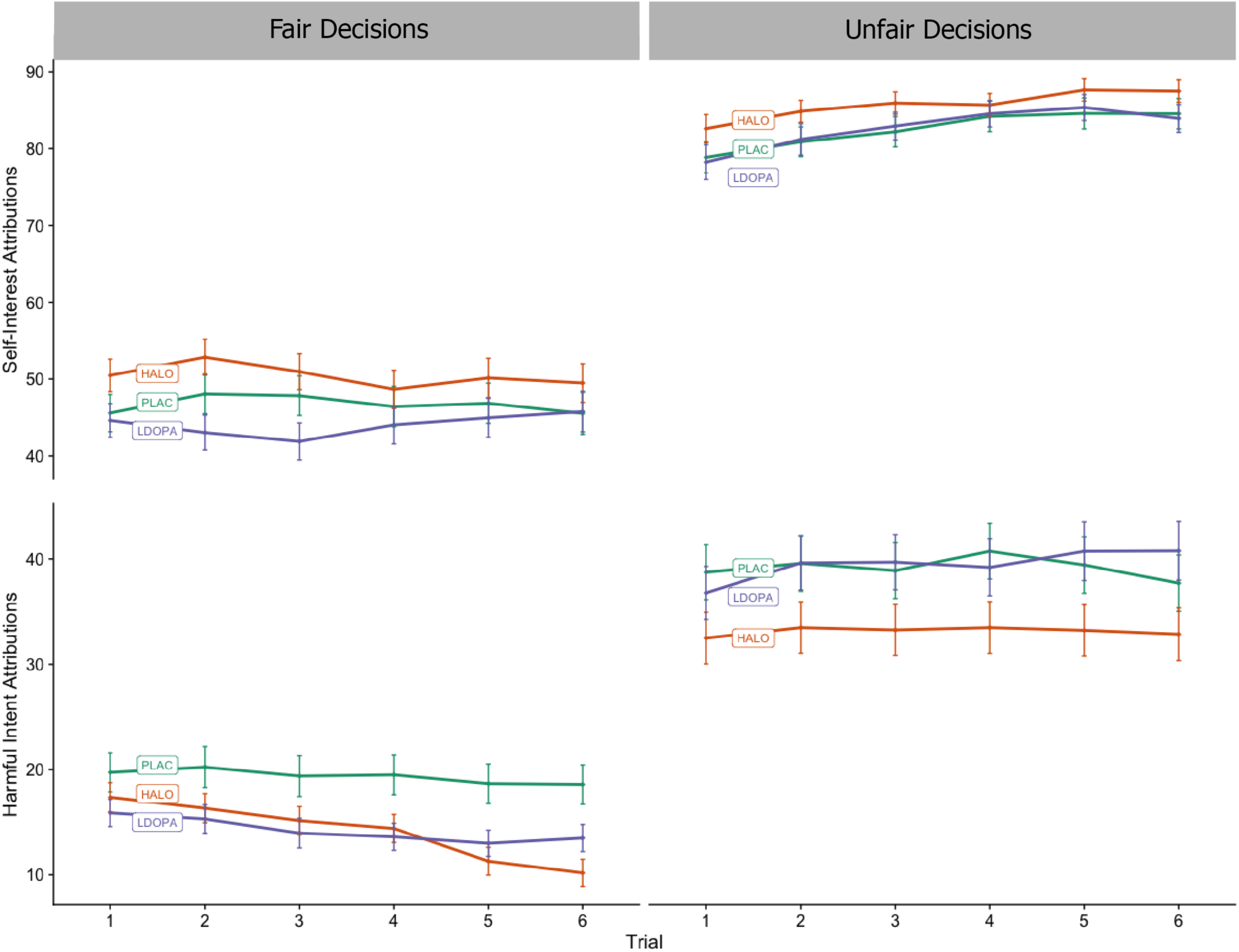
Trial by trial mean attributions of participants playing the multi-round Dictator Game for each drug condition, faceted by dictator type. Bars are the standard error of the mean. Partners were presented randomly to participants. For each trial, partners decided whether to split or keep £0.10; in unfair conditions, they always chose to keep it, and for fair conditions they always chose to split it. After each decision, participants attributed on a scale of 0-100 how much they thought their partner wanted to increase their own bonus (self-interest) and how much they thought their partner wanted to reduce their bonus (harmful intent). Relative to placebo, haloperidol demonstrates a reduction in harmful intent attributions across all dictator conditions. In fair conditions, haloperidol also demonstrates an increase in self-interest attributions. Relative to placebo, L-DOPA demonstrates a decrease of harmful intent attributions in fair conditions, and no difference compared to placebo in unfair conditions.

We also conducted an additional analysis including drug condition, trait paranoia (GPTS Total), session number, dictator, and age, with ID and Trial as random effects.

For the main effect of drug condition across conditions, detailed in Table 2 (and illustrated in Appendix E), haloperidol reduced harmful intent attributions versus placebo, but increased self-interest attributions. L-DOPA showed no effects versus placebo (although we note marginal non-significance – the upper confidence interval at zero – for harmful intent attributions versus placebo with a small effect). Haloperidol decreased harmful intent but increased self-interest attributions versus L-DOPA. The unfairness of dictators and session number both increased harmful intent attributions (Table 2).

Total GPTS summed score did not have an effect on harmful intent nor self-interest attributions. However, previous work (Bird et al., 2017) has instead used the Persecutory Ideation subscale of the GPTS as a term to assess paranoia, and so we also ran a model with this subscale as a term instead of the GPTS total. In this model, Persecutory Ideation was associated with an increase in harmful intent attribution but not self-interest attribution.

### 2.3 Post hoc analysis of changes in subjective mood and scepticism with attributions

We calculated whether there were any subjective effects of the drug on task performance by associating the change in the alertness subscale and tranquillity subscale (Herbert et al., 1976) between pre and post-dose, and then additionally between drug and placebo conditions, with harmful intent and self-interest attributions. We found that mood changes were not associated with harmful intent or self-interest attributions (p’s > 0.05; see Supplementary Material for plot). Likewise, we calculated whether participants’ beliefs about whether they were playing a real person influenced their harmful intent or self-interest attributions in any dictator condition under placebo, L-DOPA, or haloperidol. Participants were required to rate how much they believed they were playing against a real person on a scale of one to five, from ‘very sceptical’ to ‘totally believed the person was real’. At session one (first time being exposed to the game), 24 participants scored three or over, at session two, 20 participants score three or over, at session three, 21 participants scored three or over. We found that scepticism did not correlate with harmful intent or self-interest attributions for any drug or dictator condition (p’s > 0.05, see Supplementary Material).

**Table 2:** Top model average for Harmful Intent Attributions and Self-Interest attributions by drug, dictator, session number, paranoia, and age. ID and Trial Number were included as fixed effects. Model parameters: Harmful Intent/Self Interest Attributions ∼ Drug + Dictator + Paranoia + Session Number + Age + (1|ID) + (1|Trial). Models were selected and averaged based on their AICc criterion automatically in the “MuMIn” package. Beta estimates indicate the relationship between a term and harmful intent/self-interest attributions. GPTS Total was included in the model, however we also report here the post-hoc statistics of the same model with the Persecutory Ideation subscale as a term instead. We also report the difference between L-DOPA, and haloperidol run in a separate model, as our main model only compared each active condition to placebo.

| Parameter | Estimate | Standard Error | 95% CI |  | Relative Importance |
| --- | --- | --- | --- | --- | --- |
|  |  |  | Lower | Upper |  |
| Harmful Intent Attributions |  |  |  |  |  |
| Intercept 1\|2 | 2.45 | 2.09 | -1.64 | 6.54 |  |
| Intercept 2\|3 | 3.91 | 2.09 | -0.18 | 8.00 |  |
| Intercept 3\|4 | 5.44 | 2.09 | 1.34 | 9.53 |  |
| Intercept 4\|5 | 6.17 | 2.09 | 2.07 | 10.26 |  |
| Drug<br>(Haloperidol vs. Placebo) | -0.61 | 0.09 | -0.78 | -0.44 | 1 |
| Drug<br>(L-DOPA vs. Placebo) | -0.16 | 0.08 | -0.33 | 0.00 | 1 |
| Drug<br>(L-DOPA vs Haloperidol) | 0.45 | 0.09 | 0.27 | 0.62 | 1 |
| Dictator<br>(Fair < Partially Fair < Unfair) | 1.60 | 0.06 | 1.47 | 1.73 | 1 |
| Paranoia (GPTS Total) | 0.08 | 0.29 | -0.65 | 1.42 | 0.22 |
| Paranoia (Persecutory) | 1.14 | 0.45 | 0.26 | 2.02 | - |
| Session Number (1 < 2 < 3) | 0.19 | 0.06 | 0.07 | 0.31 | 1 |
| Age | 0.07 | 0.07 | -0.01 | 0.22 | 0.68 |
| Self Interest Attributions |  |  |  |  |  |
| Intercept 1\|2 | -6.10 | 1.16 | -8.38 | -3.82 |  |
| Intercept 2\|3 | -5.11 | 1.16 | -7.39 | -2.83 |  |
| Intercept 3\|4 | -3.79 | 1.16 | -6.07 | -1.52 |  |
| Intercept 4\|5 | -2.28 | 1.16 | -4.55 | 0.00 |  |
| Drug<br>(haloperidol vs. Placebo) | 0.43 | 0.08 | 0.27 | 0.59 | 1 |
| Drug<br>(L-DOPA vs. Placebo) | -0.10 | 0.08 | -0.25 | 0.05 | 1 |
| Drug<br>(L-DOPA vs haloperidol) | -0.53 | 0.08 | -0.69 | -0.37 | 1 |
| Dictator<br>(Fair < Partially Fair < Unfair) | 2.44 | 0.07 | 2.30 | 2.57 | 1 |
| Paranoia (GPTS Total) | -0.19 | 0.33 | -0.84 | 0.46 | 0.3 |
| Paranoia (Persecutory) | -0.28 | 0.34 | -1.09 | 0.10 | 0.57 |
| Session Number (1 < 2 < 3) | -0.34 | 0.06 | -0.45 | -0.23 | 1 |
| Age | -0.10 | 0.04 | -0.17 | -0.03 | 1 |

### 2.4 Multi-round Dictator Game Prediction 2: Dopamine manipulation will increase the rate at which high harmful intent attributions are reached, but not self-interest attributions

We conducted four preregistered analyses. There was no difference between L-DOPA and placebo (−0.37, 95% CI: −0.79, 0.05) or haloperidol and placebo (0.01, 95% CI: −0.45, 0.46) conditions for the trial where a harmful intent attribution score over 60 was triggered for unfair dictators. This was also true when running post hoc analysis using the relative mean of the population for each dictator (haloperidol vs placebo: 0.28, 95% CI: −0.12, 0.69; L-DOPA vs placebo: 0.16, 95% CI: −0.24, 0.55).

Because so few people scored above 60 in any trial with fair partners before the final trial our model was unable to converge. We therefore ran a post-hoc analysis with the threshold set as the mean of the population (15.87). There was no difference between L-DOPA and placebo (0.01, 95% CI: −0.33, 0.32) or haloperidol and placebo (0.15, 95% CI: −0.19, 0.49) conditions for the trial where a harmful intent attribution score over 15.87 was triggered for fair dictators.

There was no difference between L-DOPA and placebo (0.01, 95% CI: −0.25, 0.26) or haloperidol and placebo (0.05, 95% CI: −0.20, 0.30) conditions for the trial where a self-interest attribution score over 60 was triggered for unfair dictators. There was no difference between L-DOPA and placebo (0.09, 95% CI: −0.26, 0.45) or haloperidol and placebo (0.00, 95% CI: −0.35, 0.35) conditions for the trial where a self-interest attribution score over 60 was triggered for fair dictators.

### 2.5 Post-hoc analysis of change in score over time

To quantify whether dopamine manipulation adjusted the change in scores over trials for each dictator, we conducted a paired, within-subject analysis to assess the change in attributions between trial 1 and trial 6 under each drug condition.

Only haloperidol compared to placebo during unfair dictators demonstrated a reduction in harmful intent attributions between trials 1 and 6 (t (26) = 3.68, p = <0.001). There were no differences between drug conditions to changes in self-interest attributions between trial 1 and 6 (See Supplementary Material for plot).

### 2.5 Beliefs and Values Inventory

We administered the Beliefs and Values Inventory (Barnby et al., 2019b) each study day after dosing. We predicted that manipulating dopamine would moderate the ratings of interest and self-relevance of paranormal beliefs.

We found that versus placebo, neither L-DOPA nor haloperidol changed the ratings of interest or self-relevance of paranormal statements. In exploratory analyses, we found that versus placebo, neither haloperidol nor L-DOPA changed any other dimensions of agreement, self-relevance or interest across themes of science, morality, politics, and religion (see Supplementary Material).

## 3.0 Discussion

We conducted a within-subjects, double-blind, randomised controlled study examining the effects of pharmacological manipulation of the dopamine system on attributions and beliefs in healthy participants. We found that modulating dopamine led to changes in social attributions relevant to paranoia but not the salience of beliefs across multiple themes. As predicted, and consistently across conditions, haloperidol reduced attributions of harmful intent versus placebo for opponents’ actions in a multi-round Dictator Game. Additionally, against predictions, haloperidol increased self-interest attributions against placebo. In contrast, L-DOPA showed no difference versus placebo for attributions of harmful intent, except in the fair condition where they were reduced. L-DOPA versus placebo reduced self-interest attributions in fair and unfair, but not partially fair conditions. Against predictions, we found that neither haloperidol nor L-DOPA influenced the rate at which attributions of increased harmful intent were made during serial interactions. As expected, Dictator fairness and pre-existing persecutory ideation both increased attributions of harmful intent, even when taking into account drug condition, replicating previous findings and providing evidence for the validity of the paradigm.

Our results were unlikely to be a general effect of sedation or reduction in social sensitivity, as haloperidol either had absent or condition-dependent opposite effects on measures of self-interest attribution for the same events. This suggests an important selective role for dopamine in attributions of harmful intent.

Current models of antipsychotic drug action propose that blockade of post-synaptic D2 receptors in striatal regions reduces aberrant salience thereby reducing psychotic symptoms (Howes & Kapur, 2009). While therapeutic effects in patients with psychosis are generally thought to take from days to weeks to establish, the present results suggest that D2 blockade is also associated with acute reductions in attributions of harmful intent in healthy individuals. This is consistent with proposals that D2 blockade produces acute effects on cognition (Mehta et al. 1999). While we cannot be certain of the brain regions underpinning our observed effects on social cognition, it is notable that striatal D2 receptors are associated with treatment effects of D2 antagonists in psychosis (Kaar et al. 2019), although dopaminergic agents provide important modulatory function in the prefrontal cortex.

While results from this study suggest that the dopamine system is likely to have a direct role in social attributions and particularly those relevant to paranoia, current mechanistic models of the role of dopamine in psychosis cite perceptual and cognitive factors that poorly account for its social content (Bell et al., 2018). This may largely be because most experimental work on humans has focused on its role in general, rather than social cognition - for example, non-social reward (Pessiglione et al., 2006), risk and decision-making (Rutledge et al., 2015), or non-social belief updating (Nour et al., 2018). Given that we report evidence for the role of dopamine in appraisal of social threat, we suggest that dopamine modulates state representations of the social environment, much as non-social representations (e.g. stimulus reward relationships) are encoded by the interplay between the striatum and prefrontal cortex (Gershman & Uchida, 2019; Niv, 2019). Indeed, it has been previously suggested that the integration of information in the striatum is critical for social interactions and relationships (Baez-Mendoza & Schultz, 2013). Specifically, we suggest that dopamine may modulate the representation of threat during social interactions, as social threat is an evolutionarily important focus of attention (Raihani & Bell, 2019). Evidence from mice, for example, suggests a specific subcortical dopaminergic circuit for environmental threat detection and avoidance (Menegas et al., 2018). The present findings in healthy participants indicate involvement of dopamine in attributions of harm; this may be relevant for attributions subsequently incorporated into normative or pathological beliefs (Deeley 2019).

An unpredicted finding was that alongside decreasing attributions of harmful intent, haloperidol increased attributions of self-interest. This may indicate a more general involvement of dopamine in judgments about whether the intentions of social agents relate to the self or others. For example, reductions in attributions that behaviour is motivated by harmful intent may add inferential weight to alternative or competing appraisals of intention – such that disadvantageous behaviour is motivated by self-interest. However, L-DOPA was not associated with overall changes to attributions of self-interest indicating that the influence of dopamine manipulations on self-interest within this study is not symmetrical. This may be related to different mechanisms of action of the two compounds, with haloperidol blocking neurotransmission via post-synaptic dopamine D2 receptors and L-DOPA potentiating presynaptic dopamine synthesis.

We speculate that context dependent effects of L-DOPA (an effect limited to the fair condition) may reflect an interaction between the drug and the salience of others’ behaviours. We might have expected potentiating dopamine to increase paranoid attributions from the aberrant salience model (Howes & Kapur, 2009), although this model does not specify potential distinctions between paranoid attributions and those driven by presumed self-interest. Instead, we found that L-DOPA reduced attributions of both self-interest and harmful-intent under fair conditions only. There are three key models of dopamine and behaviour within which we can frame these findings. While we do not have direct measures of dopamine activity, these models warrant consideration and may provide explanatory value (while not being mutually exclusive).

First, our findings may be explained by the sigmoidal model of dopamine, where dopamine increases a neuronal population’s response to strong inputs while diminishing it for weak inputs (Servan-Schreiber et al., 1990). This fits with the L-DOPA findings for observed attributions of harmful intent and self-interest if we assume fair behaviour by a dictator provides a ‘weak’ input, and the unfair behaviour provides a stronger input (but still insufficient to be increased by L-DOPA). However, this model would predict increased attribution of harmful intent for haloperidol in the fair condition, whereas haloperidol decreased attributions of harmful intent.

The second model is a signal-to-noise account of dopaminergic modulation of neuronal activity and behaviour. Dopamine manipulations are known to affect signal-to-noise ratio, with L-DOPA predicted to both increase phasic signals while simultaneously increasing post-synaptic signal detection thresholds via increased tonic levels of dopamine (Durstewitz & Seamans, 2008; Grace, 2001). Indeed, prior experimental evidence suggests that administration of L-DOPA in healthy, sceptical individuals reduces perceptual sensitivity, with the authors suggesting this was better explained by L-DOPA *decreasing* rather than increasing signal:noise ratio (Krummenacher et al., 2010). This model also requires the assumption that social behaviours produce different input signals at different levels of fairness. In this framework, under fair conditions, the input signal may be too weak to overcome a higher set threshold for attributing intentions to another agents’ behaviour (fitting the observed reduction in self-interest *and* harmful intent). By contrast, unfair conditions are more salient and therefore readily cross a higher set threshold for attributing intentions that would otherwise be made without L-DOPA (in the placebo condition). Conversely, there is a reduction in overall signalling via post-synaptic D2 receptor blockade with haloperidol. This may explain the reduction in harmful-intent attributions, although does not easily explain the increase in self-interest attributions. Changes in attributions of self-interest may be better understood by the reductions in attributions of harmful intent adding inferential weight to the alternative/competing appraisals of intention.

Finally, the effect of dopamine manipulations are often interpreted in an ‘inverted-U’ model, whereby increases or decreases in dopamine outside an optimal signalling window lead to a decrease in behavioural response. Non-linear effects of dopamine modulation have been reported in decision-making (van der Schaaf et al., 2011), working memory (Cools & D’Esposito, 2011), sensation-seeking (Gjedde et al., 2010), and lexical decision tasks (Krummenacher et al., 2010). The data presented here suggest this may also be extended to social cognitive function in general and attributions of harmful-intent and self-interest, specifically. Within this inverted-U model, haloperidol reduced attributions of harmful-intent by reducing overall post-synaptic dopamine transmission via D2 receptors to the left of the optimum. At the same time self-interest attributions increase suggesting a separate inverted-U model for different attributions. Increased DA transmission can disrupt behaviour (Cools & D’Esposito 2011), and for this model to fit with our L-DOPA findings would also require the added assumption that different task conditions likely have different optimal dopamine levels (Zahrt et al. 1997). Thus, L-DOPA may raise dopamine release above optimal levels in fair conditions, but not potentiate dopamine enough outside optimal levels in partially fair or unfair conditions to make a difference to attributions – where a different optimal level and inverted-U model may apply.

Another possibility is the lack of significant increase in harmful-intent attributions with the administration of L-DOPA overall may be attributed to the dose being insufficient. However, we find this less likely, as L-DOPA affected other aspects of the task and prior studies using L-DOPA at the same dose showed modulation of decision-making processes, including those made within a social context (Crockett et al., 2015; Rutledge et al., 2015).

Other factors may explain the findings we observed for L-DOPA. The lack of a significant increase in harmful intent attributions under unfair conditions with L-DOPA may reflect a ceiling effect in participants with low levels of trait paranoia. It may also be that persistent increases in presynaptic dopamine release over time, coupled with sustained environmental stresses (including threatening behaviours), leads to sustained increases in attributions of harmful intent as the basis for paranoid beliefs. Paranoid states produced by drugs such as amphetamine, typically happen at high doses or after persistent use (Lecomte et al., 2018), and it has been suggested that this also occurs with the use of L-DOPA in Parkinson’s disease (Ffytche et al., 2017).

We also did not find an effect of either dopamine manipulation on the salience of paranormal beliefs – selected as an analogue of delusional ideation – and assessed using the BVI (Barnby et al. 2019b) alongside beliefs about politics, morality, religion, and science. Aberrant salience models of psychosis (e.g. Kapur et al., 2005) suggest that delusional beliefs are the outcome of sustained disruptions to striatal dopamine. Consequently, it may be that relatively brief changes to dopamine transmission are not sufficient to produce detectable changes in the salience of propositional beliefs, for which attitudes tend be more stable (Barnby et al. 2019b).

### Limitations

We use a relatively short social inference task that may preclude assessment of behaviours over a longer period of time. Previous non-social tasks (e.g. Pessiglione et al. 2006) and more recent studies with iterative social interactions (e.g. Diaconescu et al., 2017), have used comparatively longer trial designs, some in excess of 100 trials. It could be argued that some dynamics of social inference may not be evident without viewing more decisions. It remains an open question as to whether our observed drug effects would be sustained given longer social interactions, or whether we may observe sensitised responses. Also, we only used one dose of each compound and additional doses could potentially reveal a non-linear dose-response profile. There are some obvious sampling biases in our design, namely that we use all males, have a relatively small sample, and have recruited healthy individuals that happened to see our advertisement from the local community.

## Conclusions

We conducted a double-blind, within-subjects randomised controlled study in healthy individuals to test the effect of dopamine modulation on social inferences related to paranoia. We report evidence for the role of dopamine in the attribution of others’ intent to harm. Importantly, our findings were not attributable to subjective mood, beliefs in general, nor scepticism about whether participants were playing real partners. These findings are consistent with imaging and physiological evidence (Baez-Mendoza et al., 2013), and evolutionary accounts (Raihani & Bell, 2019), that identify a key role for dopamine in social inference. Future research should aim to use live, social process-oriented tasks in combination with imaging and pharmacology to better understand the role of dopamine in social attributions and its interaction with psychosocial factors (such as social stress) which are known to increase risk for psychosis.

## 4.0 Methods

### 4.1 Ethics and recruitment

This study was approved by KCL ethics board (HR-16/17-0603). All data were collected between August 2018 and August 2019. Participants were recruited through adverts in the local area, adverts on social media, in addition to adverts circulated via internal emails.

86 participants were preliminarily phone screened. 35 participants were given a full medical screen. 30 healthy males were recruited to take part in the full procedure. Inclusion criteria were that participants were healthy males, between the ages of 18 and 55. Participants were excluded if they had any evidence or history of clinically significant medical or psychiatric illness; if their use of prescription or non-prescription drugs was deemed unsuitable by the medical team; if they had any condition that may have inhibited drug absorption (e.g. gastrectomy), a history of harmful alcohol or drug use determined by clinical interview, use of tobacco or nicotine containing products in excess of the equivalent of 5 cigarettes per day, a positive urine drug screen, or were unwilling or unable to comply with the lifestyle guidelines. Participants were excluded who, in the opinion of the medical team and investigator, had any medical or psychological condition, or social circumstance which would impair their ability to participate reliably in the study, or who may increase the risk to themselves or others by participating. Some of these criteria were determined through telephone check for non-sensitive information (age, gender, general understanding of the study, overall health) before their full screening visit.

### 4.2 Procedure

Participants attended four days in total at the Centre for Neuroimaging Sciences in Denmark Hill. The first day consisted of the full medical screen that lasted approximately an hour. Participants were excluded if they were currently unwell (e.g. a cold), or if they had begun any new medication that was deemed unsuitable by medical staff. Participants underwent urinalysis, a drug screen (testing for Amphetamines, Barbiturates, Benzodiazepines, Cocaine, THC, Methadone, Methamphetamine, Opiates, Phencyclidine, and Tricyclic Antidepressants); participants were rejected if they tested positive for any of the above. Participants were also weighed and measured, and any participants with a BMI over 30 were excluded.

Electrocardiograms were taken, and participants excluded if parameters were exceeded (QTc: 330-430; PR: 120-210; QT: 270-470; QRS: <120; Heart Rate: 40-90 bpm). Additionally, blood pressure was taken, with acceptable mmHg within 90-140 (systolic) and 40-90 (diastolic) when supine and after 2 mins of standing.

A neurological assessment was made by the medical team, testing for tremor, nystagmus, pupillary reactivity, reflex test, finger-nose test, Romberg’s sign, gait, shoulder girdle strength, upper extremities strength, lower extremities strength, and myoclonic jerks. General appearance, dermatological signs, respiratory signs, cardiovascular health, abdominal signs, extremities, and musculoskeletal signs were all assessed, and participants included if normal.

Participants were given a full psychiatric exam by the medical team and excluded if any clinically significant signs or symptoms were reported, either currently or historically.

Participants then completed the OCEAN personality questionnaire (John & Srivastava, 1999), Brief O-LIFE (Mason et al., 2005), Green Paranoid Thoughts Scale (Green et al., 2008), and Bond and Lader mood rating scale (Bond & Lader, 1974).

At least 7 days later participants were then invited back for the first study session if they had satisfactorily passed the assessment day. Participants were paid £20 if they failed the screening day. Each study day was spaced by at least 7 days, but no more than two months. Each study day was identical in procedure. Participants were requested to abstain from alcohol and caffeine at least 24 hours before the study day. Study days began with a similar screening procedure to the screening day. ECGs, blood pressure, urinalysis, drug screening, neurological and physical checks were all completed upon arrival. Participants were also asked to complete the Bond and Lader mood rating scale prior to initial dosing.

Participants were initially dosed in the morning between 9.30 and 10.30am. Participants were randomly (in a Williams Square design; Williams, 1949) administered 3mg of haloperidol in two capsules or placebo in two capsules, and 10mg of Domperidone or placebo in one capsule (three capsules total).

After an hour and a half, participants were dosed a second time. This would randomly be assigned as 150mg of co-beneldopa or placebo in two capsules. Participants never took both haloperidol and co-beneldopa on the same day. Participants were also provided with a light lunch following the second dosing session. Participants only drank water throughout the entirety of the day.

In sum, participants were either given Haloperidol (3mg) + Placebo, Domperidone (10mg) + L-DOPA (150mg), or Placebo + Placebo.

After an hour and a half, participants were dosed a second time. They would randomly be assigned 150mg of co-beneldopa or placebo in two capsules. Participants never took both haloperidol and co-beneldopa on the same day. Participants were also provided with a light lunch following the second dosing session. Participants only drank water throughout the entirety of the day.

Participants were then discharged. Discharge consisted of an ECG, blood pressure assessment, neurological, and physical exam by the medical team. If participants required a taxi they were provided with one. If participants reported any adverse events these were recorded.

### 4.3 The multi-round Dictator Game

We developed a within-subjects, multi-trial modification on the Dictator game design used in previous studies to assess paranoia (Barnby et al., 2019a). Each participant played six trials against three different types of dictator. In each trial, participants were told that they have been endowed with a total of £0.10 and their partner (the dictator) had the choice to take half (£0.05) or all (£0.10) the money from the participant. Dictator decisions were one of three types: either to always take half of the money, have a 50:50 chance to take half or all of the money, or always take all of the money, labelled as Fair, Partially Fair, and Unfair, respectively. The order that participants were matched with dictators was randomised. Each dictator had a corresponding cartoon avatar with a neutral expression to support the perception that each of the six trials was with the same partner.

After each trial, participants were asked to rate on a scale of 1-100 (initialised at 50) to what degree they believed that the dictator was motivated a) by a desire to earn more (self-Interest) and b) by a desire to reduce their bonus in the trial (harmful intent). Following each block of 6 trials participants were asked to rate the character of the dictator overall by scoring intention again on both scales. Therefore, participants judged their perceived intention of the dictator on both a trial-by-trial and partner level.

After making all 42 attributions (two trial attributions for each of the 6 trials over 3 partners, plus three additional overall attributions for each partner), participants were put in the role of the dictator for 6 trials – whether to make a fair or unfair split of £0.10. Participants were first asked to choose an avatar from nine different cartoon faces before deciding on their 6 different splits. These dictator decisions were not used for analysis but were collected in the first phase of the game to match subsequent participants with decisions from real partners.

The modification to the original dictator game design allowed us to track how partner behaviour, order of partner, and whether attributions were highly variable or consistent as pre-existing paranoia changed. All participants were paid for their completion of the GPTS, regardless of follow up. Participants were paid a baseline payment for their completion.

### 4.4 Analysis

We used an information-theoretic approach for all analyses unless otherwise stated. Following Barnby et al. (2019a), we analysed the data using multi-model selection with model averaging (Burnham & Anderson, 2004; Grueber et al., 2011). The Akaike information criterion, corrected for small sample sizes (AICc), was used to evaluate models, with lower AICc values indicating a better fit (Grueber et al., 2011). The best models are those with the lowest AICc value. To adjust for the intrinsic uncertainty over which model is the true ‘best’ model, we averaged over the models in the top model set to generate model-averaged effect sizes and confidence intervals (Burnham & Anderson, 2004). In addition, parameter estimates, and confidence intervals are provided with the full global model to robustly report a variable’s effect in a model (Galipaud et al., 2014). This used package “MuMIn” (Barton, 2018). All analyses were conducted in R (Team R & R Development Core Team, 2016). All visualisations were generated using the package ‘ggplot2’ (Wickham, 2016).

In our models, all baseline continuous scale scores were centred and scaled to produce Z values. All model statistics reported are beta coefficients.

Scores of harmful intention attributions and self-interest for each dictator were taken over six trials for analysis. These were used for cumulative link mixed models (clmm; Christensen et al., 2015). In our confirmatory analysis, for each dictator harmful intent or self-interest attributions were set as our dependent variables and ID set as a random term:

Formula: Value (Ordinal) ∼ Drug + (1|ID)

In our exploratory analysis, harmful intent and self-interest attributions were set as our dependent variable. Paranoia (GPTS and Persecutory subscale), dictator behaviour (fair, unfair, partially fair), age, drug (Placebo/haloperidol/L-DOPA) were set as our explanatory terms with ID and Trial set as random terms.

Formula: Value (Ordinal) ∼ Drug + Paranoia + Dictator + Session Number + Age + (1|ID) + (1|Trial)

For our second prediction, participants that scored above 60 were considered to have scored high harmful intent attributions. Both harmful intent and self-interest scores participants were set a value of 6 if they had scored 60 in their first trial, 5 if they had scored over 60 by their second trial, 4 if they had scored 60 by their third trial, and so on. All trials following the threshold being reached were coded as 0. Participants not reaching the threshold for any trial were coded 0 across all trials. Both unfair and fair dictator behaviour were analysed with two cumulative link mixed models (clmm) each, one for harm-intent and one for self-interest.

Formula: Trial (where score > 60 triggered) ∼ Drug + (1|ID)

For attribution changes between trials one and six for each dictator and attribution type we used the R package “ggstatsplot” (Patil, 2018).

## Acknowledgements

Thanks to Dr Ndaba Mazibuko, Steph Stephenson, Dr Hannah Memon, Dr Pierluigi Selvaggi, Dr Robert McCutcheon, and Dr Stephen Kaar for being important medical support on the study. Thanks to Kirsten Brown for collecting part of the pilot data not reported in this study and assisting with ethical approval. Special thanks to Dr Uri Hertz for giving us permission to use his avatar images for the multi-round Dictator task.

## Contributions

MM & QD initially conceived of the study. JMB conceived and developed the social element of the experiment. JMB recruited participants. JMB collected the data. JMB analysed the data. JMB wrote the initial draft of the manuscript. JMB, VB, QD and MM critically revised the manuscript.

## Supplementary Material

**Appendix A.**
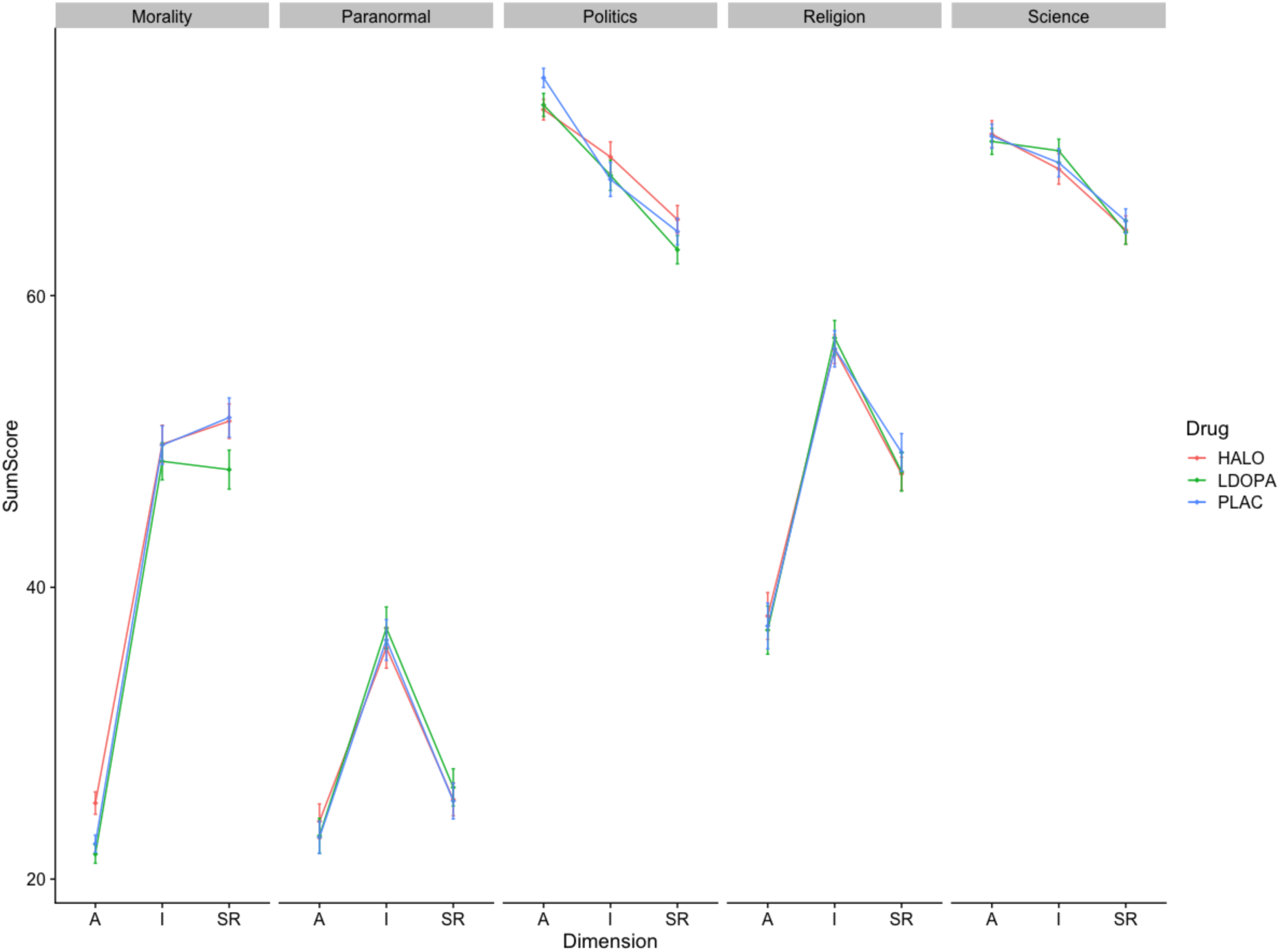
The Beliefs and Values Inventory across conditions. Aggregate scores of participants that answered the Beliefs and Values Inventory in each drug condition. Scores are divided by themes (facets) and dimensions (x-axis). Dots resemble the mean. Bars represent the standard error of the mean.

**Appendix B.**
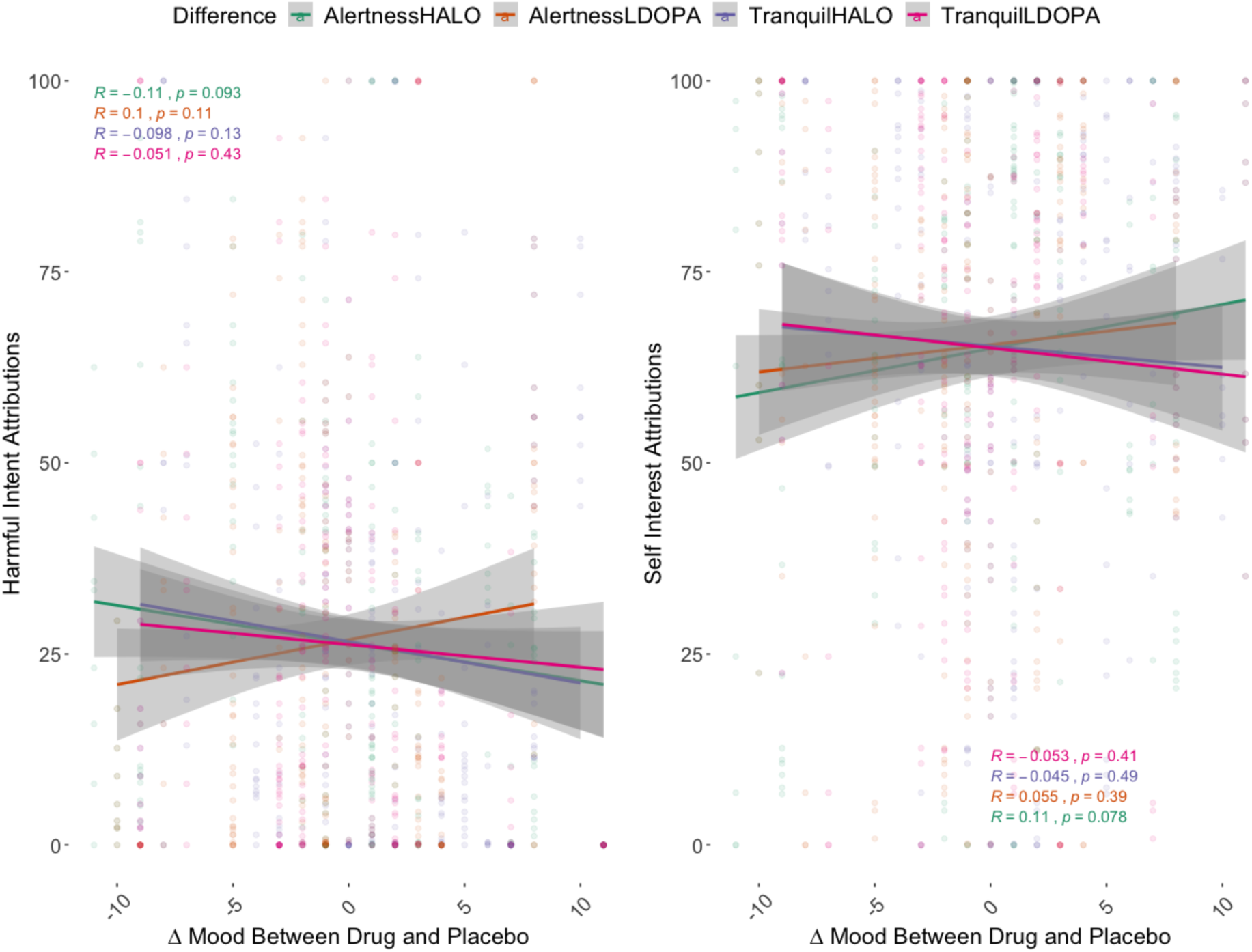
Changes in subjective mood between drug and placebo conditions following dosing. Aggregated association of scores on the Alertness and Tranquil subscales of the Bond and Lader Visual Analogue Scale between LDOPA and Placebo, and haloperidol and Placebo conditions, with Harmful Intent and Self Interest attributions. Grey shading represents the standard error of the mean.

**Appendix C.**
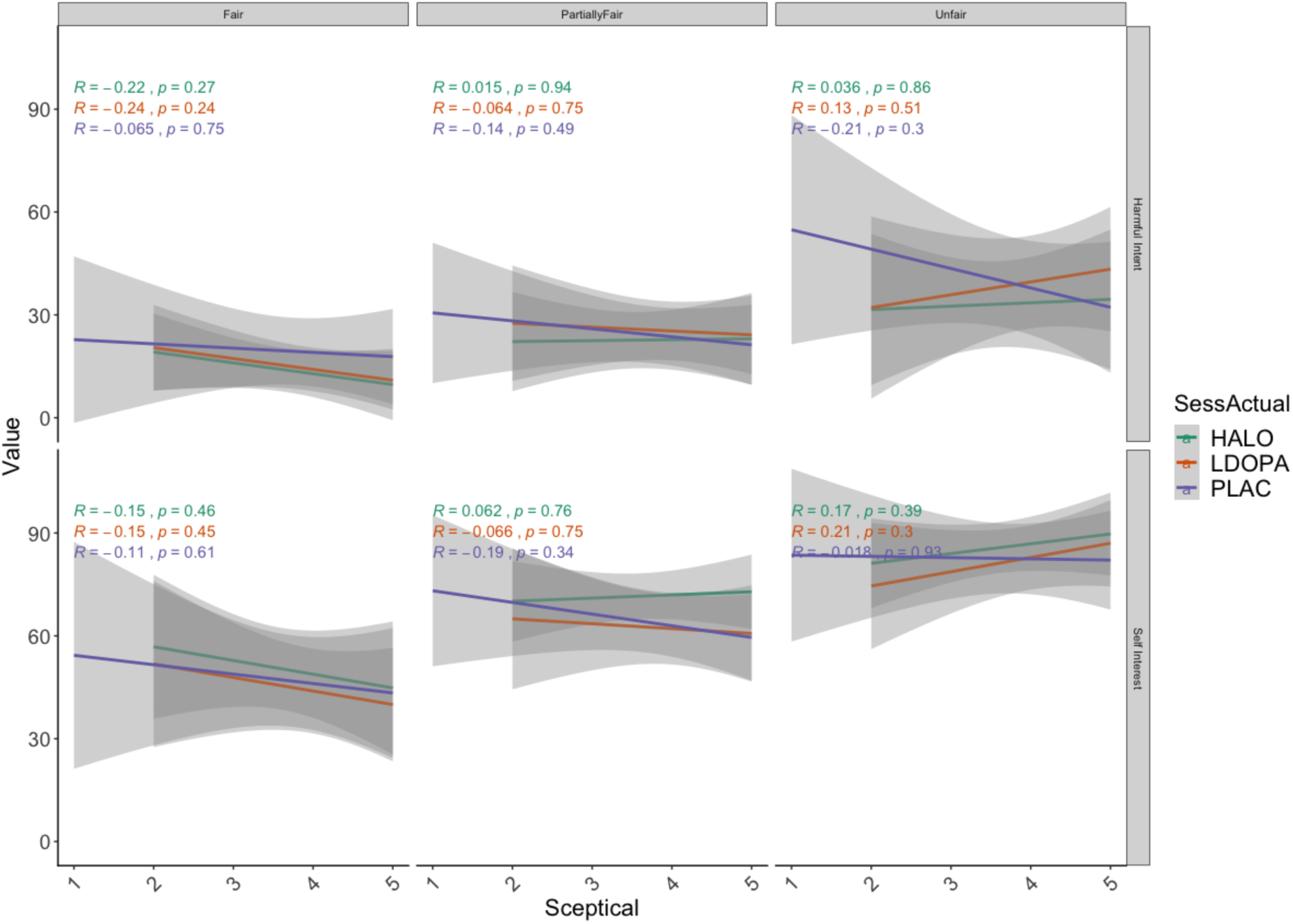
Association between scepticism scores and attributions across dictator conditions. Grey shading represents the standard error of the mean.

**Appendix D.**
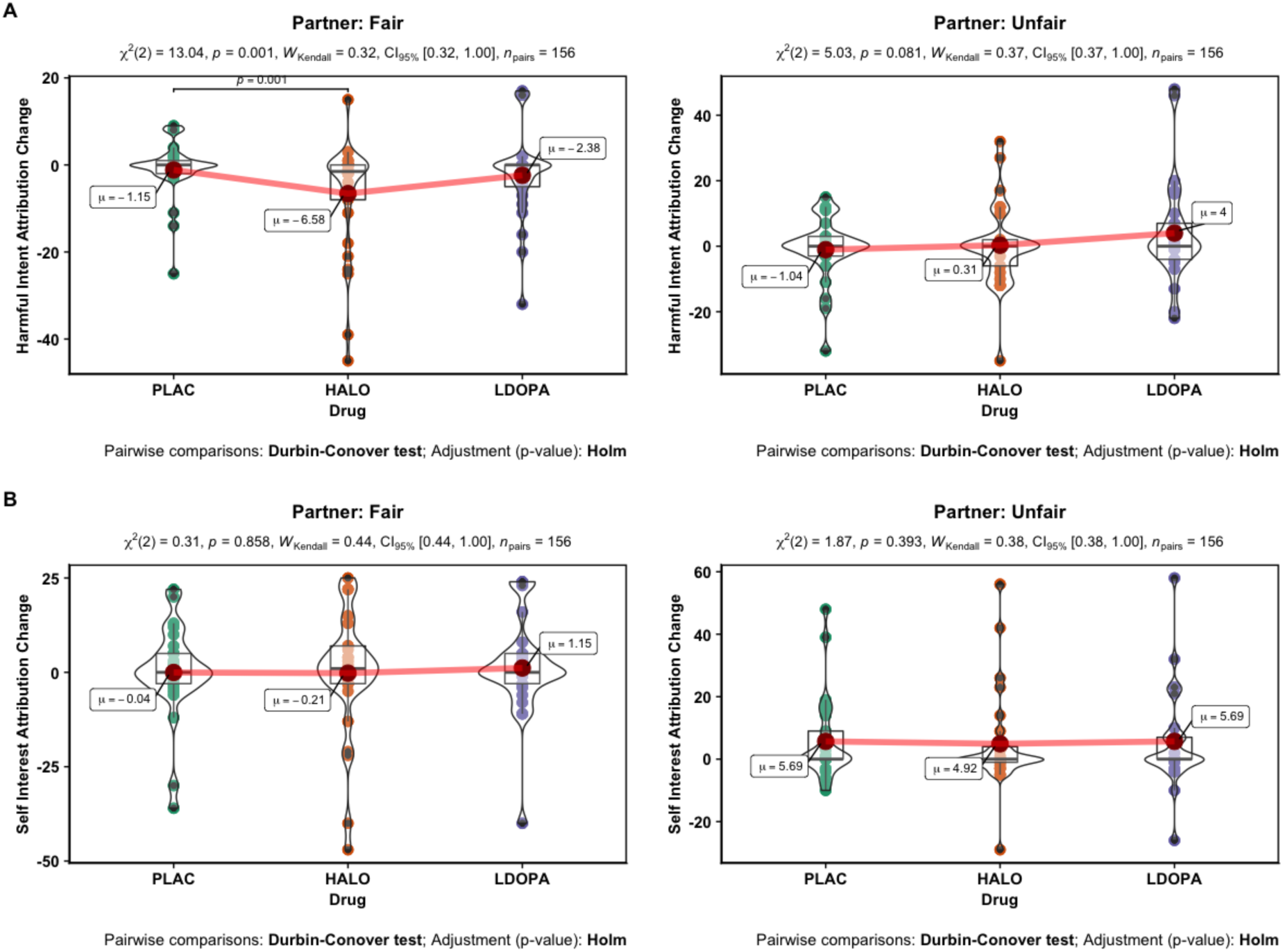
Change in attribution scores between trial 1 and 6 for unfair and fair partners for each drug condition.

**Appendix E.**
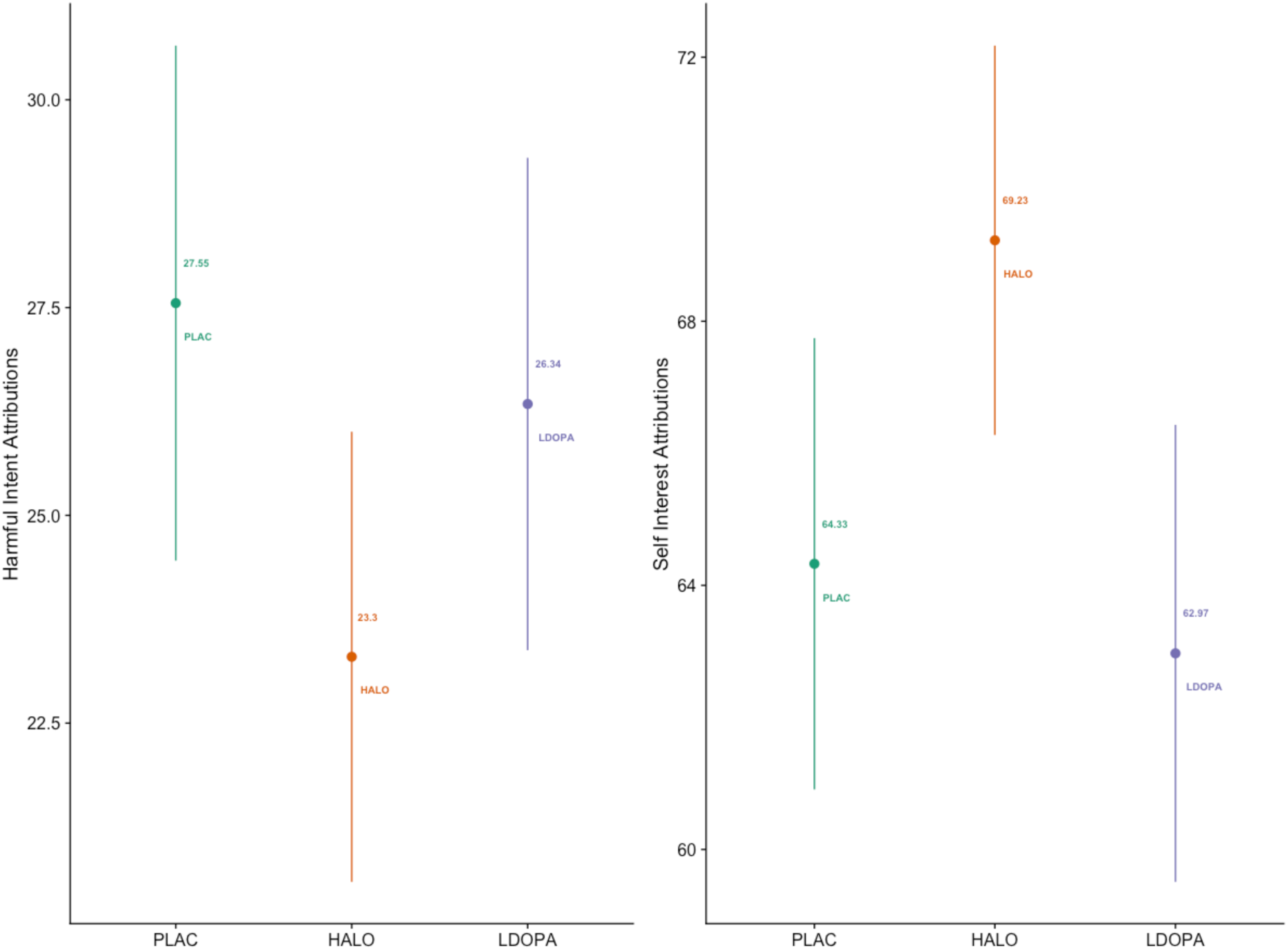
Mean and Standard Deviation of Harmful Intent Attributions and Self Interest Attribution for each drug condition, collapsed across social conditions and trials (unfair, fair, and partially fair).

## References

Andrade, L. H., & Wang, Y. P. (2012). Prevalence of psychotic symptoms in the general population varies across 52 countries. Evidence-based mental health, 15(4), 105.

Báez-Mendoza, R., & Schultz, W. (2013). The role of the striatum in social behavior. Frontiers in Neuroscience, 7, 233.

Barnby, J. M., Deeley, Q., Robinson, O. J., Raihani, N., Bell, V., & Mehta, M. (2019a). Paranoia, sensitisation and social inference: findings from two large-scale, multi-round behavioural experiments. PsyArXiv. DOI: 10.31234/osf.io/2j7ms

Barnby, J. M., Bell, V., Rains, L. S., Mehta, M. A., & Deeley, Q. (2019b). Beliefs are multidimensional and vary in stability over time-psychometric properties of the Beliefs and Values Inventory (BVI). PeerJ, 7, e6819.

Barton, K. (2018). Package “MuMIn” Title Multi-Model Inference. Retrieved from https://cran.r-project.org/web/packages/MuMIn/MuMIn.pdf

Bates, D., Maechler, M., Bolker, B., Walker, S., Christensen, R. H. B., Singmann, H., … & Grothendieck, G. (2011). Package ‘lme4’. Linear mixed-effects models using S4 classes. R package version, 1-1.

Bebbington, P. E., McBride, O., Steel, C., Kuipers, E., Radovanovic, M., Brugha, T., … & Freeman, D. (2013). The structure of paranoia in the general population. The British Journal of Psychiatry, 202(6), 419–427.

Bell, V., & O’Driscoll, C. (2018). The network structure of paranoia in the general population. Social psychiatry and psychiatric epidemiology, 53(7), 737–744.

Bird, J.C., Waite, F., Rowsell, E., Fergusson, E.C. & Freeman, D. (2017). Cognitive, affective, and social factors maintaining paranoia in adolescents with mental health problems: a longitudinal study. Psychiatry research, 257, 34-39.

Bell, V., Raihani, N., & Wilkinson, S. (2019). De-Rationalising Delusions. PsyArXiv. DOI: 10.31234/osf.io/4p9zs

Bond A, Lader M. (1974). The use of analogue scales in rating subjective feelings. British Journal of Medical Psychology, 47:211–218

Bronstein, M. V., Everaert, J., Castro, A., Joormann, J., & Cannon, T. D. (2019). Pathways to paranoia: Analytic thinking and belief flexibility. Behaviour research and therapy, 113, 18–24.

Burnham, K. P., & Anderson, D. R. (2004). Multimodel inference: Understanding AIC and BIC in model selection. Sociological Methods and Research. https://doi.org/10.1177/0049124104268644

Christensen, M. R. H. B. (2015). Package ‘ordinal’. Stand, 19, 2016.

Cools, R., & D’Esposito, M. (2011). Inverted-U–shaped dopamine actions on human working memory and cognitive control. Biological psychiatry, 69(12), e113–e125.

Corlett, P. R., Frith, C. D., & Fletcher, P. C. (2009). From drugs to deprivation: a Bayesian framework for understanding models of psychosis. Psychopharmacology, 206(4), 515–530.

Crockett, M. J., Siegel, J. Z., Kurth-Nelson, Z., Ousdal, O. T., Story, G., Frieband, C., … & Dolan, R. J. (2015). Dissociable effects of serotonin and dopamine on the valuation of harm in moral decision making. Current Biology, 25(14), 1852–1859.

Crush, E., Arseneault, L., Moffitt, T. E., Danese, A., Caspi, A., Jaffee, S. R., … & Fisher, H. L. (2018). Protective factors for psychotic experiences amongst adolescents exposed to multiple forms of victimization. Journal of psychiatric research, 104, 32–38.

Deeley, Q. (2019). Witchcraft and Psychosis: Perspectives from Psychopathology and Cultural Neuroscience. Magic, Ritual, and Witchcraft, 14(1), 86–113.

Diaconescu, A. O., Mathys, C., Weber, L. A., Kasper, L., Mauer, J., & Stephan, K. E. (2017). Hierarchical prediction errors in midbrain and septum during social learning. Social cognitive and affective neuroscience, 12(4), 618–634.

Diaconescu, A. O., Hauke, D. J., & Borgwardt, S. (2019). Models of persecutory delusions: a mechanistic insight into the early stages of psychosis. Molecular psychiatry, 1.

Durstewitz, D., & Seamans, J. K. (2008). The dual-state theory of prefrontal cortex dopamine function with relevance to catechol-o-methyltransferase genotypes and schizophrenia. Biological psychiatry, 64(9), 739–749.

Egerton, A., Chaddock, C. A., Winton-Brown, T. T., Bloomfield, M. A., Bhattacharyya, S., Allen, P., … & Howes, O. D. (2013). Presynaptic striatal dopamine dysfunction in people at ultra-high risk for psychosis: findings in a second cohort. Biological psychiatry, 74(2), 106–112.

Eisenegger, C., Pedroni, A., Rieskamp, J., Zehnder, C., Ebstein, R., Fehr, E., & Knoch, D. (2013). DAT1 polymorphism determines L-DOPA effects on learning about others’ prosociality. PLoS One, 8(7), e67820.

Esslinger, C., Englisch, S., Inta, D., Rausch, F., Schirmbeck, F., Mier, D., … & Zink, M. (2012). Ventral striatal activation during attribution of stimulus saliency and reward anticipation is correlated in unmedicated first episode schizophrenia patients. Schizophrenia research, 140(1-3), 114–121.

Floel, A., Garraux, G., Xu, B., Breitenstein, C., Knecht, S., Herscovitch, P., & Cohen, L. G. (2008). Levodopa increases memory encoding and dopamine release in the striatum in the elderly. Neurobiology of aging, 29(2), 267–279.

Freeman, D. (2007). Suspicious minds: the psychology of persecutory delusions. Clinical psychology review, 27(4), 425–457.

Freeman, D., McManus, S., Brugha, T., Meltzer, H., Jenkins, R., & Bebbington, P. (2011). Concomitants of paranoia in the general population. Psychological medicine, 41(5), 923–936.

Freeman, D., Stahl, D., McManus, S., Meltzer, H., Brugha, T., Wiles, N., & Bebbington, P. (2012). Insomnia, worry, anxiety and depression as predictors of the occurrence and persistence of paranoid thinking. Social psychiatry and psychiatric epidemiology, 47(8), 1195–1203.

Freeman, D., & Garety, P. (2014). Advances in understanding and treating persecutory delusions: a review. Social psychiatry and psychiatric epidemiology, 49(8), 1179–1189.

Fusar-Poli, P., & Meyer-Lindenberg, A. (2012). Striatal presynaptic dopamine in schizophrenia, Part II: meta-analysis of [18F/11C]-DOPA PET studies. Schizophrenia bulletin, 39(1), 33–42.

Ffytche, D., Creese, B., Politis, M., Chaudhuri, K. R., Weintraub, D., Ballard, C., & Aarsland, D. (2017). The psychosis spectrum in Parkinson disease. Nature Reviews Neurology, 13(2), 81.

Galipaud, M., Gillingham, M. A. F., David, M., & Dechaume-Moncharmont, F. X. (2014). Ecologists overestimate the importance of predictor variables in model averaging: A plea for cautious interpretations. Methods in Ecology and Evolution, 5(10), 983–991.

Gershman, S. J., & Uchida, N. (2019). Believing in dopamine. Nature Reviews Neuroscience, 20(11), 703–714.

Gjedde, A., Kumakura, Y., Cumming, P., Linnet, J., & Møller, A. (2010). Inverted-U-shaped correlation between dopamine receptor availability in striatum and sensation seeking. Proceedings of the National Academy of Sciences, 107(8), 3870–3875.

Grace, A. A. (2001). Psychostimulant actions on dopamine and limbic system function: relevance to the pathophysiology and treatment of ADHD. Stimulant drugs and ADHD: Basic and clinical neuroscience, 5, 134–157.

Green, C. E. L., Freeman, D., Kuipers, E., Bebbington, P., Fowler, D., Dunn, G., & Garety, P. A. (2008). Measuring ideas of persecution and social reference: The Green et al. Paranoid Thought Scales (GPTS). Psychological Medicine, 38(1), 101–111.

Grueber, C. E., Nakagawa, S., Laws, R. J., & Jamieson, I. G. (2011). Multimodel inference in ecology and evolution: Challenges and solutions. Journal of Evolutionary Biology. John Wiley & Sons, Ltd (10.1111).

Herbert, M., Johns, M. W., & Doré, C. (1976). Factor analysis of analogue scales measuring subjective feelings before and after sleep. British Journal of Medical Psychology, 49(4), 373–379.

Howes, O., Bose, S., Turkheimer, F., Valli, I., Egerton, A., Stahl, D., … & McGuire, P. (2011). Progressive increase in striatal dopamine synthesis capacity as patients develop psychosis: a PET study. Molecular psychiatry, 16(9), 885.

Howes, O. D., & Kapur, S. (2009). The dopamine hypothesis of schizophrenia: version III— the final common pathway. Schizophrenia bulletin, 35(3), 549–562.

Howes, O. D., & Murray, R. M. (2014). Schizophrenia: an integrated sociodevelopmental-cognitive model. The Lancet, 383(9929), 1677–1687.

John, O. P., & Srivastava, S. (1999). The Big-Five trait taxonomy: History, measurement, and theoretical perspectives. In L. A. Pervin & O. P. John (Eds.), Handbook of personality: Theory and research (Vol. 2, pp. 102–138). New York: Guilford Press.

Kaar, S. J., Natesan, S., McCutcheon, R., & Howes, O. D. (2019). Antipsychotics: mechanisms underlying clinical response and side-effects and novel treatment approaches based on pathophysiology. Neuropharmacology, 107704.

Kapur, S., Mizrahi, R., & Li, M. (2004). From dopamine to salience to psychosis—linking biology, pharmacology and phenomenology of psychosis. Schizophrenia research, 79(1), 59–68.

Krummenacher, P., Mohr, C., Haker, H., & Brugger, P. (2010). Dopamine, paranormal belief, and the detection of meaningful stimuli. Journal of Cognitive Neuroscience, 22(8), 1670–1681.

Lecomte, T., Dumais, A., Dugré, J. R., & Potvin, S. (2018). The prevalence of substance-induced psychotic disorder in methamphetamine misusers: A meta-analysis. Psychiatry research, 268, 189–192.

Niv, Y. (2019). Learning task-state representations. Nature neuroscience, 22(10), 1544–1553.

Nour, M. M., Dahoun, T., Schwartenbeck, P., Adams, R. A., FitzGerald, T. H., Coello, C., … & Howes, O. D. (2018). Dopaminergic basis for signalling belief updates, but not surprise, and the link to paranoia. Proceedings of the National Academy of Sciences, 115(43), E10167–E10176.

Mason, O., Linney, Y., & Claridge, G. (2005). Short scales for measuring schizotypy. Schizophrenia research, 78(2-3), 293–296.

McCutcheon, R. A., Abi-Dargham, A., & Howes, O. D. (2019). Schizophrenia, dopamine and the striatum: from biology to symptoms. Trends in Neurosciences, 42, 205–220.

Mehta, M. A., Sahakian, B. J., McKenna, P. J., & Robbins, T. W. (1999). Systemic sulpiride in young adult volunteers simulates the profile of cognitive deficits in Parkinson’s disease. Psychopharmacology, 146(2), 162–174.

Menegas, W., Akiti, K., Amo, R., Uchida, N., & Watabe-Uchida, M. (2018). Dopamine neurons projecting to the posterior striatum reinforce avoidance of threatening stimuli. Nature neuroscience, 21(10), 1421.

Mohr, C., Krummenacher, P., Landis, T., Sandor, P. S., Fathi, M., & Brugger, P. (2005). Psychometric schizotypy modulates levodopa effects on lateralized lexical decision performance. Journal of Psychiatric Research, 39(3), 241–250.

Patil, I. (2018). Ggstatsplot: ggplot2 Based Plots with Statistical Details. Retrieved from: https://CRAN.R-project.org/package=ggstatsplot. DOI: 10.5281/zenodo.2074621

Pedroni, A., Eisenegger, C., Hartmann, M. N., Fischbacher, U., & Knoch, D. (2014). Dopaminergic stimulation increases selfish behavior in the absence of punishment threat. Psychopharmacology, 231(1), 135–141.

Pessiglione, M., Seymour, B., Flandin, G., Dolan, R. J., & Frith, C. D. (2006). Dopamine-dependent prediction errors underpin reward-seeking behaviour in humans. Nature, 442(7106), 1042.

Raihani, N. J., & Bell, V. (2017a). Conflict and cooperation in paranoia: a large-scale behavioral experiment. Psychological Medicine, pp. 1–11.

Raihani, N. J., & Bell, V. (2017b). Paranoia and the social representation of others: A large-scale game theory approach. Scientific Reports, 7(1), 4544.

Raihani, N. J., & Bell, V. (2019). An evolutionary perspective on paranoia. Nature human behaviour, 3(2), 114–121.

Rutledge, R. B., Skandali, N., Dayan, P., & Dolan, R. J. (2015). Dopaminergic modulation of decision making and subjective well-being. Journal of Neuroscience, 35(27), 9811–9822.

Saalfeld, V., Ramadan, Z., Bell, V., & Raihani, N. J. (2018). Experimentally induced social threat increases paranoid thinking. Royal Society open science, 5(8), 180569.

Seeman, P. (1987). Dopamine receptors and the dopamine hypothesis of schizophrenia. Synapse, 1(2), 133–152.

Servan-Schreiber, D., Printz, H., & Cohen, J. D. (1990). A network model of catecholamine effects: gain, signal-to-noise ratio, and behavior. Science, 249(4971), 892–895.

Spitzer, M. (1995). A neurocomputational approach to delusions. Comprehensive psychiatry, 36(2), 83–105.

Startup, H., Freeman, D., & Garety, P. A. (2007). Persecutory delusions and catastrophic worry in psychosis: developing the understanding of delusion distress and persistence. Behaviour research and therapy, 45(3), 523–537.

Sterzer, P., Adams, R. A., Fletcher, P., Frith, C., Lawrie, S. M., Muckli, L., … & Corlett, P. R. (2018). The predictive coding account of psychosis. Biological psychiatry, 84(9), 634–643.

Team, R. D. C., & R Development Core Team, R. (2016). R: A Language and Environment for Statistical Computing. R Foundation for Statistical Computing, 1(2.11.1), 409. https://doi.org/10.1007/978-3-540-74686-7

Tsou, J. Y. (2012). Intervention, causal reasoning, and the neurobiology of mental disorders: Pharmacological drugs as experimental instruments. Studies in History and Philosophy of Science Part C: Studies in History and Philosophy of Biological and Biomedical Sciences, 43(2), 542–551.

van der Schaaf, M. E., Warmerdam, E., Crone, E. A., & Cools, R. (2011). Distinct linear and non-linear trajectories of reward and punishment reversal learning during development: Relevance for dopamine’s role in adolescent decision making. Developmental cognitive neuroscience, 1(4), 578–590.

Voce, A., Calabria, B., Burns, R., Castle, D., & McKetin, R. (2019). A Systematic Review of the Symptom Profile and Course of Methamphetamine-Associated Psychosis: Substance Use and Misuse. Substance use & misuse, 54(4), 549–559.

Volkow, N. D., Wang, G. J., Fowler, J. S., Telang, F., Maynard, L., Logan, J., … & Zhu, W. (2004). Evidence that methylphenidate enhances the saliency of a mathematical task by increasing dopamine in the human brain. American Journal of Psychiatry, 161(7), 1173–1180.

Wickham, H. (2016). ggplot2: Elegant Graphics for Data Analysis. New York: Spinger. Retrieved from https://cran.r-project.org/web/packages/ggplot2/citation.html

Wickham, S., Taylor, P., Shevlin, M., & Bentall, R. P. (2014). The impact of social deprivation on paranoia, hallucinations, mania and depression: the role of discrimination social support, stress and trust. PloS one, 9(8), e105140.

Williams, E. J. (1949): Experimental designs balanced for the estimation of residual effects of treatments. Australian Journal of Scientific Research, Ser. A 2, 149–168.

Zahrt, J., Taylor, J. R., Mathew, R. G., & Arnsten, A. F. (1997). Supranormal stimulation of D1 dopamine receptors in the rodent prefrontal cortex impairs spatial working memory performance. Journal of neuroscience, 17(21), 8528–8535.

